# Responses to Visual Speech in Human Posterior Superior Temporal Gyrus Examined with iEEG Deconvolution

**DOI:** 10.1101/2020.04.16.045716

**Authors:** Brian A. Metzger, John F. Magnotti, Zhengjia Wang, Elizabeth Nesbitt, Patrick J. Karas, Daniel Yoshor, Michael S. Beauchamp

## Abstract

Experimentalists studying multisensory integration compare neural responses to multisensory stimuli with responses to the component modalities presented in isolation. This procedure is problematic for multisensory speech perception since audiovisual speech and auditory-only speech are easily intelligible but visual-only speech is not. To overcome this confound, we developed intracranial encephalography (iEEG) deconvolution. Individual stimuli always contained both auditory and visual speech but jittering the onset asynchrony between modalities allowed for the time course of the unisensory responses and the interaction between them to be independently estimated. We applied this procedure to electrodes implanted in human epilepsy patients (both male and female) over the posterior superior temporal gyrus (pSTG), a brain area known to be important for speech perception. iEEG deconvolution revealed sustained, positive responses to visual-only speech and larger, phasic responses to auditory-only speech. Confirming results from scalp EEG, responses to audiovisual speech were weaker than responses to auditory-only speech, demonstrating a subadditive multisensory neural computation. Leveraging the spatial resolution of iEEG, we extended these results to show that subadditivity is most pronounced in more posterior aspects of the pSTG. Across electrodes, subadditivity correlated with visual responsiveness, supporting a model in visual speech enhances the efficiency of auditory speech processing in pSTG. The ability to separate neural processes may make iEEG deconvolution useful for studying a variety of complex cognitive and perceptual tasks.

**Significance statement:** Understanding speech is one of the most important human abilities. Speech perception uses information from both the auditory and visual modalities. It has been difficult to study neural responses to visual speech because visual-only speech is difficult or impossible to comprehend, unlike auditory-only and audiovisual speech. We used intracranial encephalography (iEEG) deconvolution to overcome this obstacle. We found that visual speech evokes a positive response in the human posterior superior temporal gyrus, enhancing the efficiency of auditory speech processing.

## Introduction

When humans communicate face-to-face, auditory information from the talker’s voice and visual information from the talker’s mouth both provide clues about speech content. A critical brain area for multisensory speech perception is the posterior superior temporal gyrus and sulcus (pSTG), the location of human auditory association cortex (Moerel et al., 2014; Leaver and Rauschecker, 2016). The belt and parabelt areas in pSTG are selective for both the complex acoustic-phonetic features that comprise auditory speech (Belin et al., 2000; Formisano et al., 2008; Mesgarani et al., 2014) and the mouth movements that comprise visual speech (Beauchamp et al., 2004; Bernstein et al., 2011; Rhone et al., 2016; Ozker et al., 2017; Zhu and Beauchamp, 2017; Ozker et al., 2018b; Rennig and Beauchamp, 2018; Beauchamp, 2019). However, the neural computations used by the pSTG to integrate auditory and visual speech features are poorly understood.

A widely used schema for understanding multisensory processing compares the amplitude of the responses to unisensory and multisensory stimuli (Stein and Stanford, 2008). If the responses to multisensory stimuli are greater than the sum of the responses to the component unisensory stimuli, the multisensory computation is termed “superadditive” suggesting the existence of facilitatory interactions between modalities. In contrast, if the multisensory responses are less than the sum of the unisensory responses, the computation is “subadditive” suggesting suppressive interactions between modalities. Multisensory responses equal to the sum of the unisensory responses are termed “additive”, reflecting little or no interaction between modalities.

While this schema was codified in responses to simple auditory beep and visual flash stimuli in anesthetized animals (Stein and Stanford, 2008) it has also been applied to human brain responses to auditory and visual speech recorded with BOLD fMRI, scalp encephalogram (EEG) and intracranial EEG (iEEG) (Besle et al., 2004; van Wassenhove et al., 2005; Besle et al., 2008; Rhone et al., 2016; Karas et al., 2019). In these studies, responses to unisensory and multisensory speech were compared under the assumption that different *sensory* responses were the main driver of neural activity. However, perception of visual-only speech (lip-reading or speech-reading) places very different *cognitive* demands on humans, as it is both difficult and inaccurate: visual-only speech is largely unintelligible (Fig. 1A)(Peelle and Sommers, 2015). This lack of intelligibility could lead to decreased neural activity in language areas which process semantics, increased activity in attentional control areas, or both, confounding assessment of the multisensory computation.

**Figure 1.**
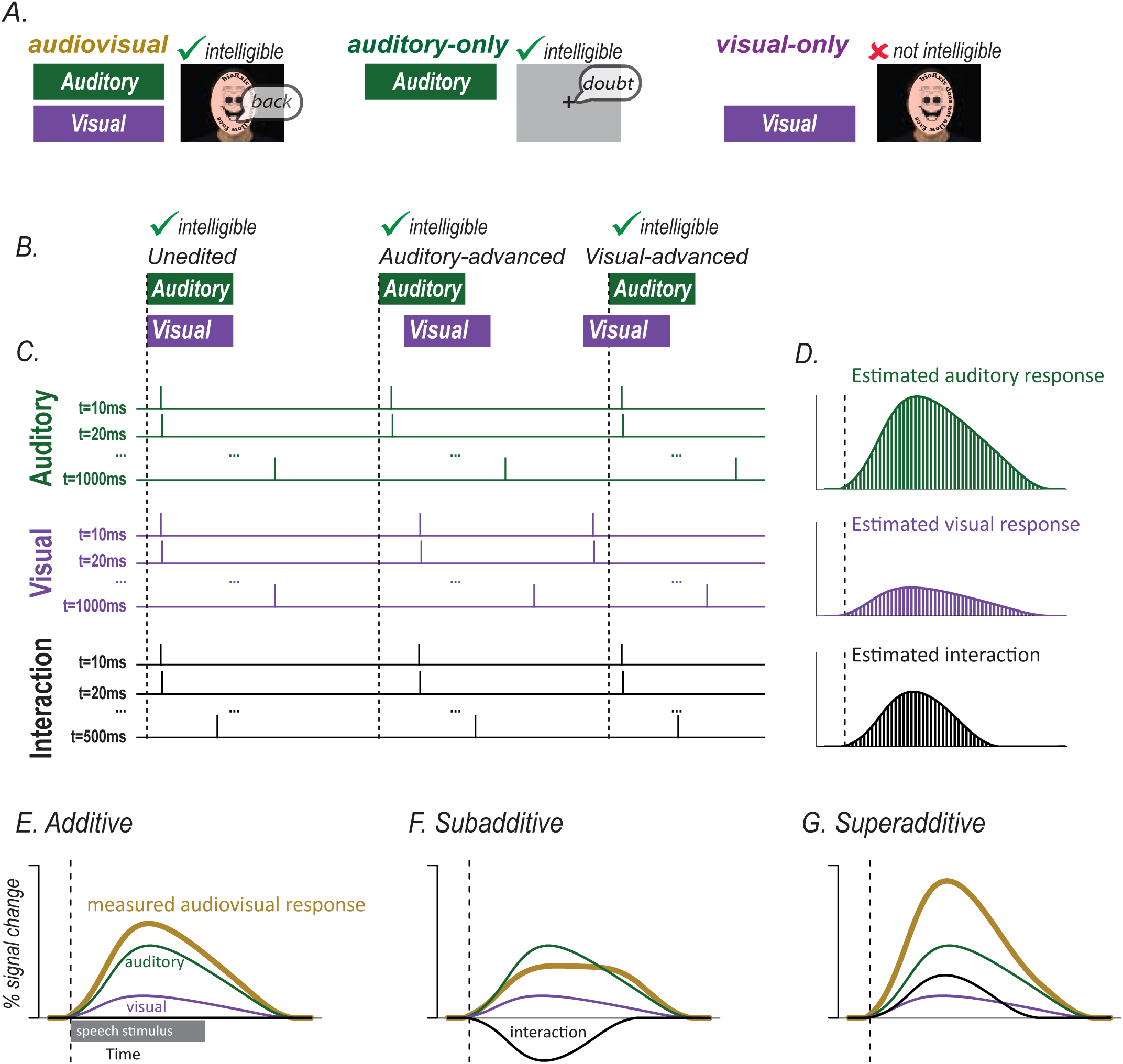
A. In many studies, audiovisual speech, auditory-only speech, and visual-only speech are presented. Comparing neural responses to these stimuli is confounded by the fact that audiovisual and auditory-only speech is intelligible while visual-only speech is not. B. An alternative approach is to present audiovisual speech with varying asynchrony. Audiovisual speech can be unedited (synchronous), edited so that the auditory component of the speech is moved earlier (auditory-advanced) or edited so that the visual component of the speech is moved earlier (visual-advanced). All three types of speech are intelligible. C. Modifying the synchrony of the auditory and visual speech components allowed for the component responses to be estimated using deconvolution. The responses to auditory and visual speech were estimated with one predictor for each 10 ms time point of the response. Each predictor consists of a delta or tent function with amplitude of 1 at a single time point of the response and an amplitude of 0 at other time points (for simplicity, only the first, second and last predictors are shown; ellipsis indicates the remainder of the predictors). The sum of the predictors were fit to the measured neural response using a generalized linear model to create an independent estimate of the auditory and visual responses. If auditory and visual speech begin at the same time, as shown for the unedited stimulus, the Audt=10 ms and Vist=10 ms regressors are identical, making it impossible to determine their relative amplitudes. If additional stimuli are presented for which the auditory and visual speech are temporally asynchronous, as shown for the auditory-advanced and visual-advanced stimuli, then the Audt=10 ms and Vist=10 ms regressors are offset. If many stimuli are presented with different asynchronies, the complete time courses of the auditory and visual responses can be estimated. The auditory and visual responses were modelled for 1 second after speech onset, the interaction response was modelled for 500 ms after speech onset. D. After the amplitude of each regressor was estimated by fitting to the measured neural response to all stimuli, the value of each time point regressor was plotted to show the time course of the component responses. The bars show the amplitude of the regressor modeling each individual time point of the response (green for auditory, purple for visual, black for interaction). E. The gold curve shows the measured response to audiovisual speech. If there are little or no interactions between modalities, the measured response will be equal to the sum of the estimated auditory (green) and visual (purple) responses. The interaction term will be zero (black line along x-axis). F. If the interactions between modalities are subadditive, the measured response will be less than the sum of the estimated auditory (green) and visual (purple) responses. The interaction term will be negative. G. If the interactions between modalities are superadditive, the measured response will be less than the sum of the estimated auditory (green) and visual (purple) responses. The interaction term will be positive.

To circumvent this problem, we applied a novel experimental design inspired by the use of finite-impulse response functions to analyze event-related BOLD fMRI (Glover, 1999). In our design, all stimuli consisted of audiovisual speech, but the temporal onset asynchrony of the auditory and visual speech was varied (Fig. 1B). Then, deconvolution was applied to estimate the component unisensory responses and the multisensory computation (Fig. 1C). Presentation of audiovisual speech with a fixed asynchrony results in summed neural responses to the auditory and visual speech components that cannot be separated due to collinearity in the resulting system of linear equations. However, if the presentation of one component is delayed, the neural response for that component is also delayed. Repeating this process for different onset asynchronies allows the system of equations to be solved for the complete time course of the unisensory responses and the interaction between them (Fig. 1D).

If the value of the interaction term is near zero, the measured audiovisual response will be equal to the sum of the auditory and visual responses, supporting an additive model of multisensory interactions during speech processing (Fig. 1E). If the interaction term is negative, the measured audiovisual response will be less than the sum of the estimated unisensory responses, indicating subadditivity (Fig. 1F). With a positive interaction term, the measured audiovisual response will be greater than the sum, indicating superadditive interactions between modalities (Fig. 1G).

The key advantage of deconvolution is that it allows for the estimation of the responses to visual and auditory speech from measurements of the response to audiovisual speech presented with varying asynchrony. The temporal binding window for auditory and visual speech is on the order of hundreds of milliseconds (Grant and Seitz, 2000; Magnotti et al., 2013; Picton, 2013; Wallace and Stevenson, 2014) perhaps due to the large variability present in natural audiovisual speech (Chandrasekaran et al., 2009; Schwartz and Savariaux, 2014). This means that audiovisual speech with varying asynchrony is readily intelligible, avoiding the confounds introduced by comparing responses to intelligible audiovisual or auditory-only speech with responses to unintelligible visual-only speech.

## Materials and methods

### Human subjects

All experiments were approved by the Committee for the Protection of Human Subjects at Baylor College of Medicine and participants provided written informed consent. Participants consisted of seven subjects (6F, mean age 37, 6L hemisphere) undergoing intracranial electrode grid placement for phase two epilepsy monitoring. Electrode grids and strips were placed based on clinical criteria for epilepsy localization and resection guidance. All experiments were conducted in the epilepsy monitoring unit and clinical monitoring continued unabated.

### Experimental Design

Visual stimuli were presented with an LCD monitor (Viewsonic VP150, 1024 x 768 pixels) placed on a stand located 57cm in front of the subject’s face. Auditory stimuli were presented through two speakers mounted on the wall behind and above the patient’s head. Stimuli were presented using the Psychtoolbox extensions for MATLAB (Brainard, 1997; Pelli, 1997; Kleiner et al., 2007).

The stimuli consisted of audiovisual recordings of four different words (“back,” “beach,” “doubt,” and “pail”) recorded at 30 Hz (video) and 44.1 kHz (audio) selected from the Hoosier Audiovisual Multitalker Database (Lachs and Hernandez, 1998; Conrey and Pisoni, 2004). Visual speech onset was defined as the time of the first video frame containing a visible mouth movement related to speech production. Auditory speech onset was defined as the first positive deflection in the auditory envelope corresponding to the beginning of the speech sound. Relative to the beginning of the recording, the visual onset/auditory onsets were: “back”, 500 ms/631 ms; “beach”, 367ms/551 ms; “doubt”, 367ms/478 ms; “pail”, 433ms/562 ms. This produced onset asynchronies of 131 ms, 184 ms, 111 ms and 129 ms. On average, the visual onset occurred 139 ms before the auditory onset.

There were three asynchrony conditions (Fig. 1B). The first condition consisted of unedited audiovisual recordings. The second condition consisted of auditory-advanced words for which the audio component was shifted forward in time by 300 ms in Adobe Premiere. The third condition consisted of visual-advanced words for which the video component was shifted forward in time by 300 ms. This resulted in twelve total stimuli (four stimulus exemplars times three asynchrony conditions). Stimuli were presented in random order and after each trial participants responded via keypress whether they perceived the audiovisual speech as synchronous or asynchronous.

### iEEG Data Collection

Neural signals were recorded with subdural grid and strip electrodes (2.3 mm exposed diameter platinum alloy discs embedded in flexible silastic sheets; Ad-Tech Corporation, Racine, WI) connected to a Cerebus data acquisition system (Blackrock Microsystems, Salt Lake City, UT). A reversed intracranial electrode facing the skull was used as a reference for recording, and all signals were amplified, filtered (high-pass 0.3 Hz first-order Butterworth, low pass 500 Hz fourth-order Butterworth) and digitized at 2000 Hz. A photodiode was placed on the stimulus monitor to capture the exact time of visual stimulus onset. Both the photodiode output and the auditory output of the stimulus presentation computer were recorded by the data acquisition system to ensure precise synchronization between sensory stimulation and the evoked neural response.

### Software tools and availability

All data analysis was conducted using the software tool RAVE (R Analysis and Visualization of intracranial Electroencephalography, freely available for Windows, Mac and Linux platforms at https://github.com/beauchamplab/rave.

### iEEG Data Analysis: Preprocessing

Data was notch filtered (60 Hz, 1st and 2nd harmonics) and converted into frequency and phase domains using a wavelet transform. The number of cycles of the wavelet was increased as a function of frequency, from 3 cycles at 2 Hz to 20 cycles at 200 Hz, to optimize tradeoff between temporal and frequency precision (Cohen, 2014). Data was down sampled to 100 Hz after the wavelet transform and then re-referenced to the average of all valid channels, determined by visual inspection. The continuous data was epoched into trials using the auditory speech onset of each stimulus as the reference (*t* = 0).

For each trial and frequency, the power data were transformed into percentage signal change from baseline, where baseline was set to the average power of the response from −1.0 to - 0.5 seconds prior to auditory speech onset. This time window consisted of the inter-trial interval, during which participants were shown a dark gray screen with a white fixation point. The percent signal change from this pre-stimulus baseline was then averaged over frequencies from 70-150 Hz to calculate the broadband high-frequency activity (BHA).

### iEEG Data Analysis: Electrode Localization and Selection

FreeSurfer (RRID:SCR_001847; (Dale et al., 1999b; Fischl et al., 1999a)) was used to construct cortical surface models for each subject from their preoperative structural T1 magnetic resonance image scans. Post-implantation CT brain scans, showing the location of the intracranial electrodes, were then aligned to the preoperative structural MRI brain using AFNI, Analysis of Functional Neuroimaging (Cox, 1996). Electrode positions were marked manually using BioImage Suite 35 (Joshi et al., 2011) and projected to the nearest location on the cortical surface using iELVis (Groppe et al., 2017). SUMA in the AFNI package was used to visualize cortical surface models with the overlaid electrodes, and positions were confirmed using intraoperative photographs of the electrode grids overlaid on the brain when available. For single subject analysis, electrodes were visualized on that subject’s cortical surface model. For group analysis, each participant’s brain was aligned to the sulcal and gyral pattern of the Colin N27 brain (Holmes et al., 1998) using the FreeSurfer spherical template (Fischl et al., 1999b). Each surface was resampled to contain exactly 198,912 nodes, with the result that a given node index had the same anatomical location in any subject (Argall et al., 2006). Then, the node index closest to each electrode was determined in each subject, and the electrodes from all subjects displayed at the appropriate node in the N27 brain.

From a total of 786 electrodes implanted in 7 patients, *n* = 33 electrodes were selected that were located in the pSTG and showed a significant neural response (31 left hemisphere, 2 right hemisphere, electrode locations shown in Figure 2A). The pSTG was defined as the portion of STG located posteriorly to the central sulcus, if the central sulcus continued in an inferior direction. A significant neural response was defined as *p* < 0.001, Bonferroni-corrected BHA response to all speech words in the window from auditory stimulus onset to stimulus offset (0 seconds to 0.5 seconds). Because the functional criterion ignored word type, the main comparisons of interest were independent of the functional criterion and hence unbiased.

**Figure 2.**
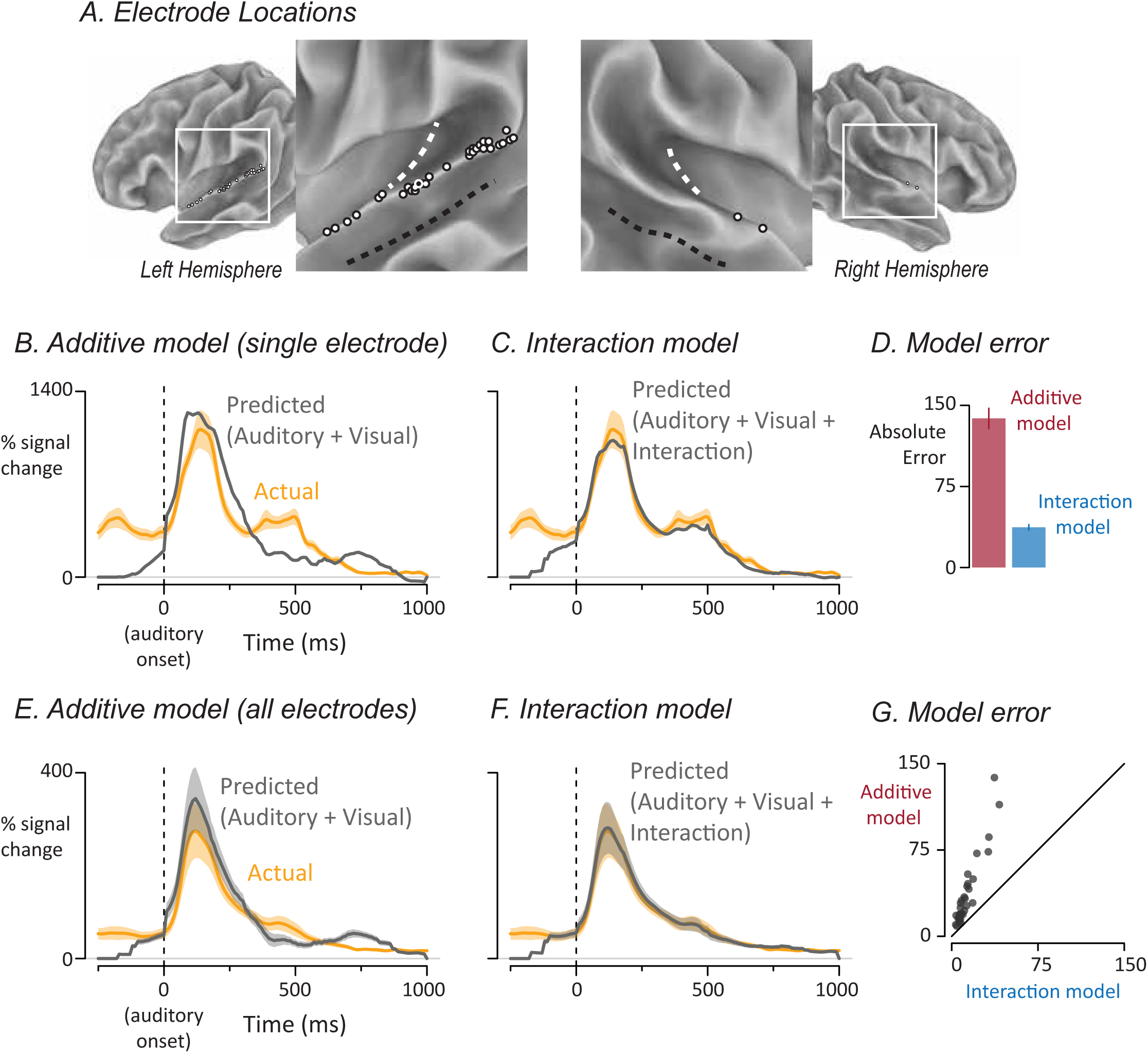
A. The location of left-hemisphere (left panel, *n* = 31) and right-hemisphere (right panel, *n* = 2) electrodes that met both an anatomical criterion (located on the posterior superior temporal gyrus) and a functional criterion (significant response to speech) displayed on a partially inflated cortical surface model. The white dashed line shows Heschl’s gyrus, the black dashed line shows the fundus of the superior temporal sulcus. Each electrode is shown as a white circle with a black outline, except for one electrode shown as a black circle with a white outline, corresponding to electrode YAR-22, responses shown in (B), (C), and (D). B. The yellow line shows the measured response of a single electrode (YAR-22) to audiovisual speech with auditory speech onset at time zero and response in units of percent increase of broadband high-frequency power from pre-stimulus baseline. The mean and standard error of the mean (SEM, yellow shaded area) at each time point was calculated across trials. The gray line shows the predicted time series from the additive model in which auditory and visual responses were summed. C. For this electrode, the predicted response (gray line) of the interaction model in which auditory, visual and interaction estimates were summed. Actual response same as in (B). D. For this electrode, the error for each model was calculated by taking the absolute difference between the predicted and measured response at each time point and calculating the mean and SEM across time points. E. The yellow curve shows the mean and SEM across electrodes of the response to audiovisual speech. For each electrode, an additive model was created in which auditory and visual responses were summed to generate a predicted audiovisual response. Gray curve shows mean and SEM of the predicted response across electrodes. F. For each electrode, an interaction model was created in which auditory, visual and interaction responses were summed to generate a predicted response. Actual response same as in (E). G. Model error for the additive and interaction models plotted against each other, one symbol per electrode.

### iEEG Data Analysis: Deconvolution

Inspired by analyses in which deconvolution was used to separate temporally overlapping responses in event-related fMRI designs (Glover, 1999), we used deconvolution to decompose the measured iEEG responses to audiovisual speech into responses to auditory and visual speech. In order to implement deconvolution in a generalized linear model, a separate regressor was used to model every time point of the response (Fig. 1C). Since the regressors are independent, allowing neural responses to be modelled without any assumption as to their shape. In the resulting fitted models, the fit coefficient (beta-weight) for each regressor can be plotted (Fig. 1D), providing a best-fit estimate of the response over time to that stimulus (similar to a time-locked average).

The first model was an additive model in which auditory and visual speech evoke time-varying responses that sum at each time point to produce the measured response. To fit the additive model, we constructed two sets of regressors to estimate the time course of the response of each electrode to the auditory and visual components of speech. The time base of the regressors was the same as that of the data (with a value every 10 ms) and the data window extended for 1 second from the onset of the given modality, resulting in 99 time points for each of the auditory and visual responses (the response at the time of stimulus onset was constrained to be zero). Each regressor consisted of a single stick function (also known as a delta function or tent function) at the appropriate post-stimulus time, and was zero everywhere else, so that fitting the entire set of regressors modeled the time course of the response without any assumptions about its shape. As shown by the design matrix in Figure 1-1, this resulted in a total of 198 different regressors, equivalent to 198 free parameters, one for each time point of the auditory and visual responses.

**Figure 1-1.**
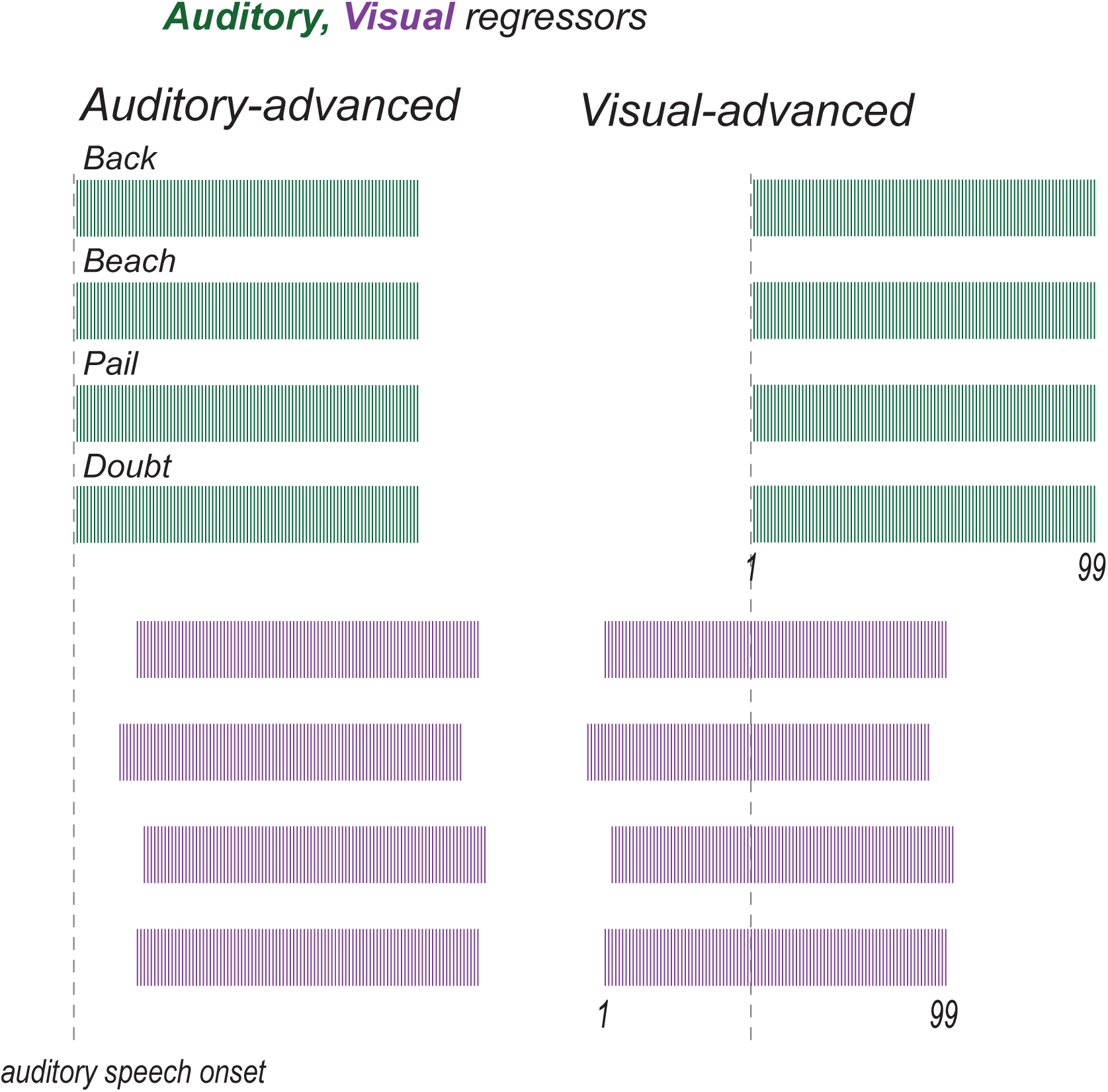
For the additive model, auditory (green) and visual (purple) regressors were fit to auditory- advanced words (left column) and visual-advanced words (right column). Unisensory responses were modeled using 99 regressors, starting 10 ms after auditory onset, spaced at 10 ms intervals. Each regressor consisted of a single tent or stick function (vertical colored line) that was “1” at the appropriate post-stimulus onset time and zero everywhere else (as shown in Fig. 1C). For efficiency, all 99 regressors are shown collapsed onto a single time axis. The auditory regressors for the four different words (”back”,”beach”,”pail”,”doubt”) were aligned to auditory speech onset (dashed grey line). The visual regressors for the four different words were adjusted to reflect the actual onset time of visual speech for each word.

In event-related fMRI, a single stimulus condition usually consists of presentation of multiple exemplars (for instance, a face, followed a few seconds later by a different face image, and so on); the deconvolved impulse response function represent the average response across all faces even though it is understood that individual stimuli evoke differing responses (Westfall et al., 2016). Our implementation of iEEG deconvolution is equivalent to this approach: the deconvolved visual, auditory, and interaction responses represent the average response across the four different word stimuli. An alternative approach would be to create separate deconvolution regressors for each word, but this would greatly increase the number of free parameters (to 198 parameters per word * 4 words = 792 free parameters).

Responses to the four different word stimuli were aligned so that the auditory onset of each word occurred at time zero (this was necessary because the precise onset time of auditory speech within each unedited video clip varied slightly). The visual regressors were also shifted in time according to the timing of the visual speech in each clip. For instance, for the stimulus exemplar “doubt” in the visual-advanced condition, prior to fitting, the visual regressor was shifted to begin at *t* = −411 ms and end at *t* = 589 ms to reflect the exact timing of the onset of visual speech for that exemplar/condition pair.

Deconvolution relies on variable asynchrony between the events in order to prevent collinearity (Fig.1C). In rapid-event-related fMRI experiments, randomized stimulus schedules are used that introduce variable asynchrony between different trial types (Dale et al., 1999a). In our design, collinearity was addressed using the auditory-visual asynchrony in the twelve different trial types. In particular, the auditory-advanced and visual-advanced conditions had the least overlap between the auditory and visual regressors, reducing collinearity. For the additive model, ordinary least squares regression was used to determine auditory and visual coefficients for every time point in the auditory-advanced and visual-advanced conditions, and these fitted coefficients were used to predict the response to the unedited condition by summing the auditory and visual coefficients at each time point. To provide the best possible prediction, as during the fitting process, the visual coefficients were aligned based on the actual auditory-visual onset asynchrony.

The second deconvolution model included an interaction term (similar to that used in an ANOVA) in addition to the unisensory auditory and visual responses. Like the unisensory responses, the interaction term was modelled by creating a set of stick functions, with value “1” at the appropriate post-stimulus time point and zero everywhere else. While the unisensory responses were modelled for 1 second after stimulus onset, the interaction could only be modelled for 500 ms; this was the duration for which most trials had both auditory and visual speech present. The first time point of the interaction term occurred at auditory onset, and the final time point occurred at the offset of visual speech in the natural-head-start condition, 500 ms later. The interaction time course was composed of 50 different regressors, corresponding to one sample every 10 ms, and were generated by taking the product of the two unisensory regressors (A x V) with the additional constraint that the interaction could not occur more than 600 ms after visual onset. Ordinary least squares fitting was used to estimate the coefficients at each time point for the auditory, visual and interaction regressors simultaneously, across all conditions. Then, the three coefficients at each time point were summed to generate a predicted response. Figure 1-2 provides a representation of the entire design matrix of the interaction model.

The fitting procedure was repeated independently for every electrode. Stimulus was not included in the model, resulting in the simplifying assumption that all stimulus exemplars evoked the same amplitude of auditory and visual responses. However, the relative timing of the auditory and visual stimulus onsets of each word was used in the model, reducing the effect of condition and word to a single value, that of the relative asynchrony of auditory and visual speech. The fitted model was used to predict the response to each word (while the model contained only one coefficient for each regressor at each time point, it generated different predictions for each word and condition because of the audiovisual timing differences between them). To measure the goodness of fit, we compared the average (across words) predicted response with the actual average response in the unedited condition, from 100ms before speech onset (to capture visual-only activity) to 1000ms after speech onset. We calculated error by taking the absolute difference between the predicted and actual curves at each time point. Averaging error across time points provided a single measure of error for each electrode.

**Figure 1-2.**
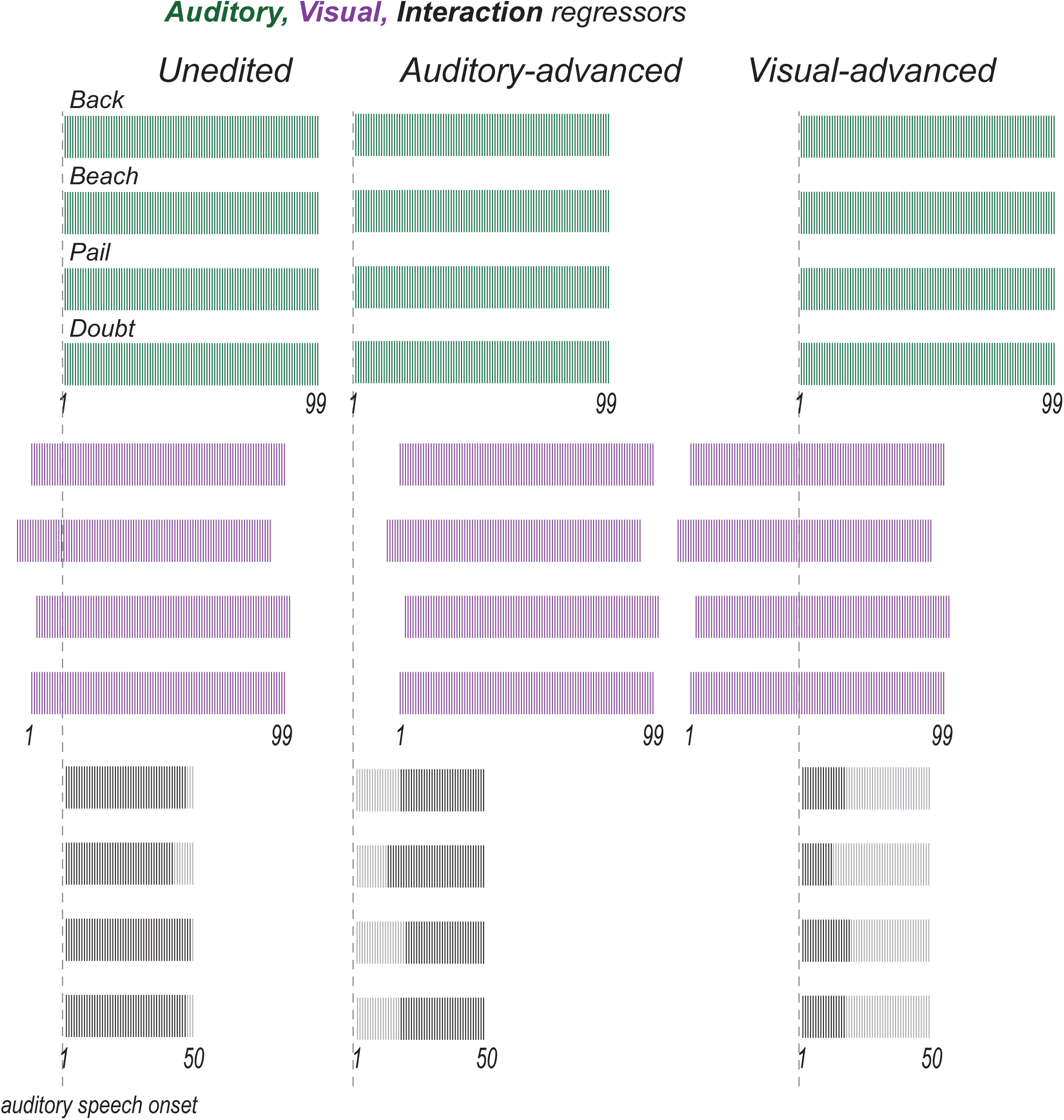
For the interaction model, auditory (green), visual (purple) and interaction (black) regressors were fit to unedited words (left column), auditory-advanced words (middle column) and visual- advanced words (right column). The auditory regressors for the four different words were aligned to auditory speech onset (dashed grey line). The visual regressors for the four different words were adjusted to reflect the actual onset time of visual speech for each word. The interaction response was modeled using 50 regressors starting 10 ms after auditory onset, spaced at 10 ms intervals. Interaction regressors were weighted 1 (black sticks) when there was an overlap between auditory and visual speech windows and 0 otherwise (gray sticks). Interaction regressors were also weighted zero for timepoints more than 600 ms after visual onset.

### Deconvolution Model Comparison

Linear mixed-effects modeling was used to compare the fits of the additive model (with only auditory and visual terms) and the interaction model (with auditory, visual and interaction terms) to the responses in each electrode (Schepers et al., 2014; Ozker et al., 2017; Ozker et al., 2018a; Karas et al., 2019; Sjerps et al., 2019). The lme4 package was used for model construction (Bates et al., 2015), followed by *t*-tests with Satterthwaite-approximated degrees of freedom (Kuznetsova et al., 2017). The dependent variable was model fit (absolute error). The fixed factor was model type (additive *vs.* interaction, with additive used as the baseline). The random factors were participant and electrode nested within participant.

**Figure 2-1.**
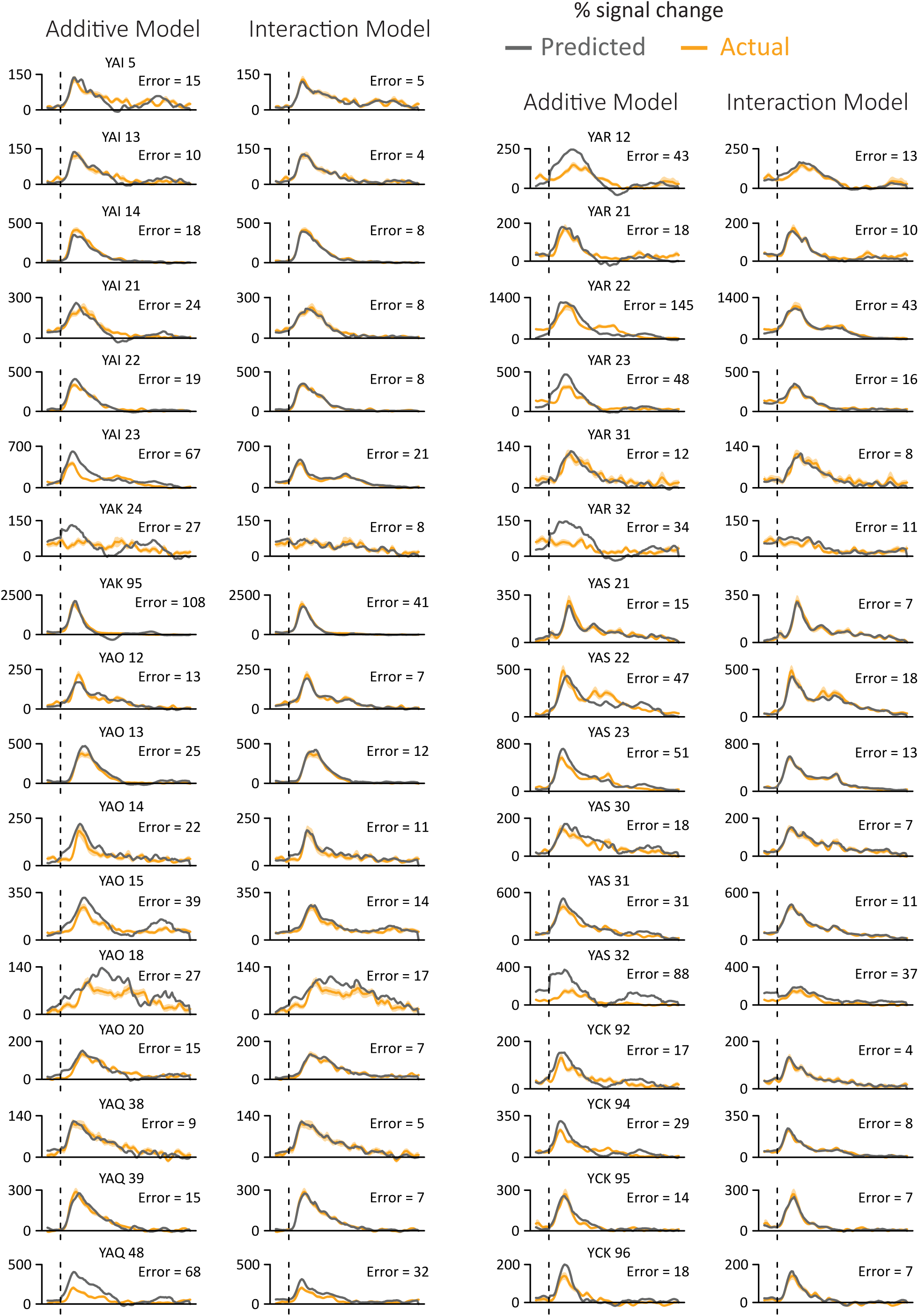
For each electrode, the yellow line shows the mean and SEM across trials of the measured response to audiovisual speech (y-axis is percent increase in BHA relative to baseline, x-axis is time, dashed line shows onset of auditory speech). In the left sub-plot, the gray line shows the predicted response from the additive model in which auditory and visual responses were summed. In the right sub-plot, the gray line shows the predicted response of the interaction model in which auditory, visual and interaction estimates were summed. Number over each sub- plot show total absolute error for that model.

### Estimates of Variance

In order to visualize the variance of the measured response to audiovisual speech in a single electrode (Fig. 2B and 2C), the response at each 10-ms timepoint of the response was measured in each trial, and the mean and standard error of the mean (SEM) at each time point across trials was calculated. The deconvolution models were fit to the data, producing a single predicted time course for the additive model and a different predicted time course for the interaction model. The average absolute error for each model was calculated by taking the absolute difference between the predicted and measured response at each time point and calculating the mean and SEM across time points (Fig. 2D).

To calculate the variance of the measured response to audiovisual speech across all electrodes, the mean and SEM at each time point across electrodes was calculated (Fig. 1E and 1F). The variance of the model fits was calculated the same way, by taking the average and SEM across electrodes at each time point (Fig. 1E and 1F). The variance of the estimated auditory, visual and interactions responses was also calculated as the mean and SEM for each time point (regressor) across electrodes (Fig. 3A).

**Figure 3.**
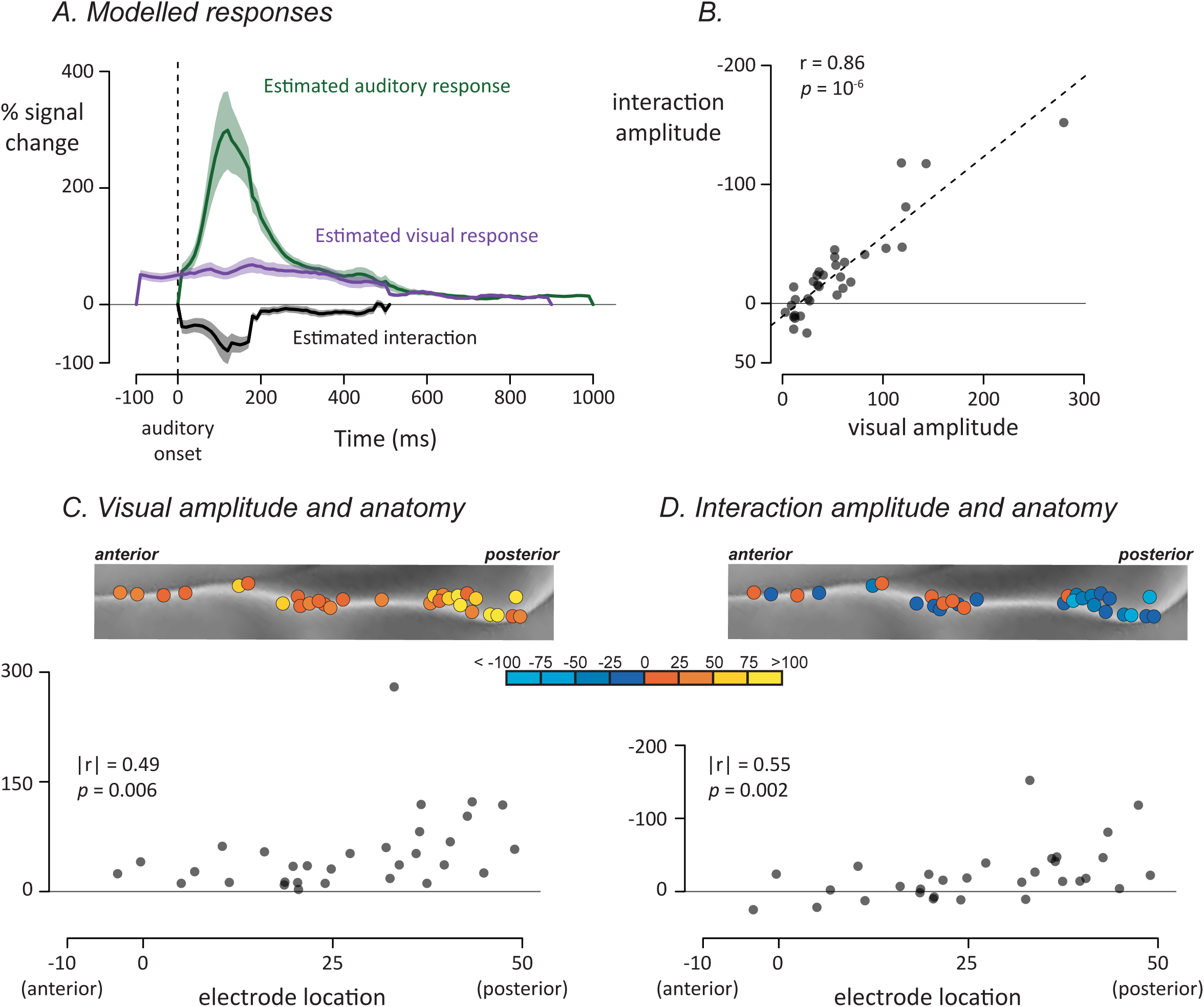
A. For each electrode, an interaction model was fit to generate modelled auditory (green), visual (purple) and interaction (black) time courses. Plots show mean and SEM across all electrodes. B. For each electrode, the visual response was plotted against the average interaction response, one symbol per electrode. C. All left hemisphere electrodes (rotated version of Fig. 2A, left panel) colored by their visual response amplitude. Bottom panel shows plot of anterior-to-posterior location in standard space against visual response amplitude. D. Left hemisphere electrodes colored by their interaction amplitude. Bottom panel shows plot of same data.

**Figure 3-1.**
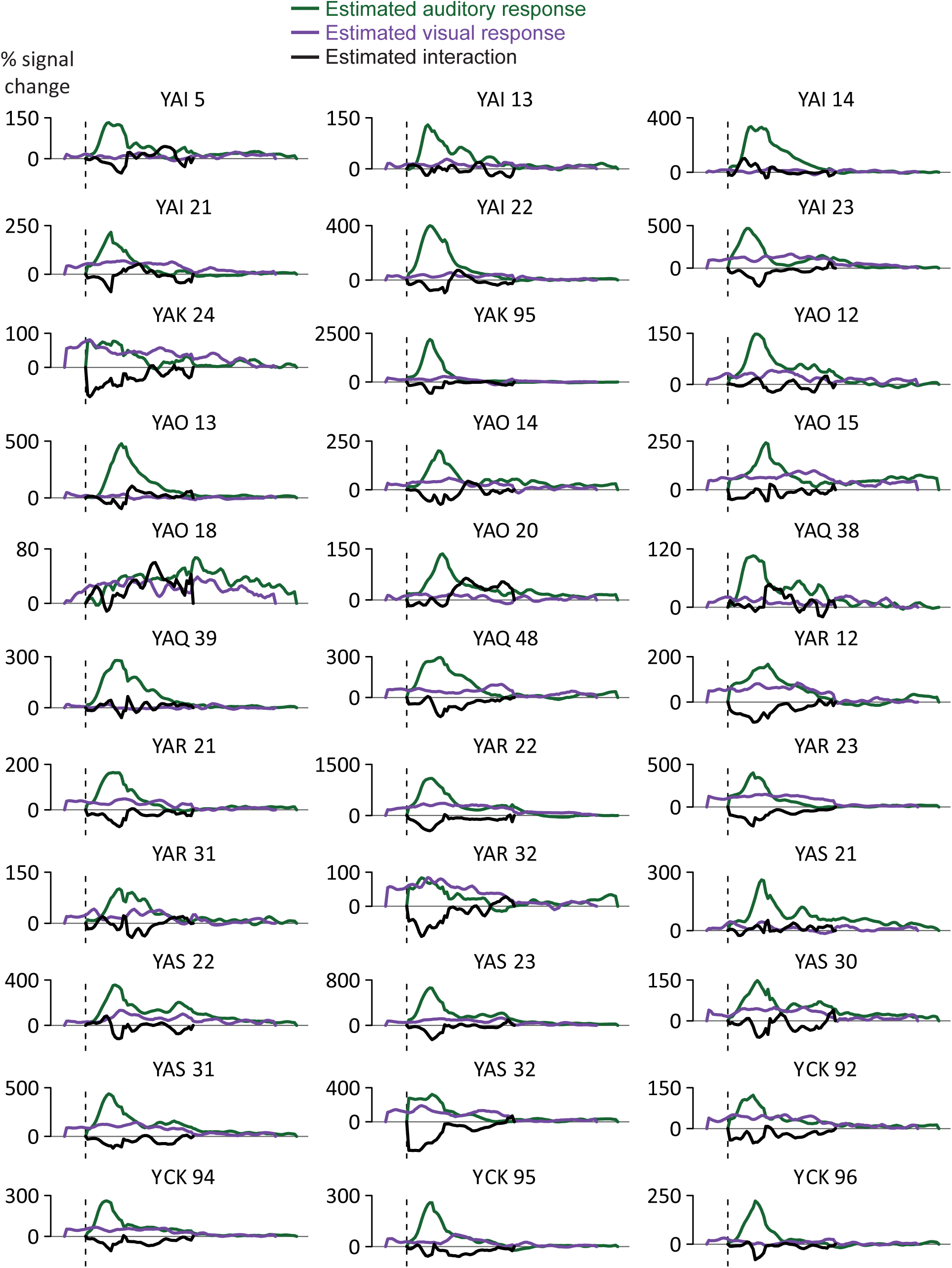
For each electrode, the green line shows the estimated auditory response, the purple line shows the estimated visual response, the black line shows the estimated interaction response (y-axis is percent increase in BHA relative to baseline, x-axis is time, dashed line shows onset of auditory speech).

### Spatial Analysis

For each electrode, an (x, y, z) coordinate was assigned from the node closest to that electrode in a cortical surface model of the N27 brain in MNI space. The y-value of the standard space coordinate provided the electrode’s location along the anterior-to-posterior axis of the pSTG (Ozker et al., 2017; Ozker et al., 2018b). To measure the correspondence between electrode location and iEEG responses, we correlated the location of the thirty-one electrodes on the left pSTG with other values (there were only two electrodes over right pSTG, too few to analyze). In some reporting conventions, more posterior anatomical locations are defined as more positive on the A-P axis, while in others, more posterior locations are defined as more negative. To avoid confusion, we report the absolute value of the spatial correlation values along with an explicit statement about the direction of the correlation (*e.g. “*greater visual response for more posterior electrodes”).

Spearman rank correlations were used in order to minimize the influence of extreme values. The electrode locations were correlated with the visual response converted to a single value by averaging the deconvolution time-point regressors over a 500 ms window beginning at the onset of the visual regressor. A single value for the interaction response was calculated by averaging the deconvolution regressors in a 500 ms window beginning at the onset of the deconvolution regressor.

To determine if these spatial patterns were consistent across participants, we also fit linear mixed-effects models. Along with the fixed effect of electrode location, the models included random intercepts and random slopes for each participant. If the spatial pattern was driven by inter-participant variability, the random slope term will capture this variance and the fixed effect of spatial location will not be statistically significant (X2 test). To make this analysis commensurate with the Spearman correlation analysis, each variable was converted to a rank value before being entered into the model.

## Results

We examined the response to audiovisual speech with varying asynchrony in 33 pSTG electrodes from 7 patients (electrode locations shown in Fig. 2A). One goal of the analysis was to determine whether interactions between neural responses to auditory and visual speech were additive, subadditive, or superadditive. The first step in making this determination was to compare two different deconvolution models, an additive model and an interaction model. Both models used as input the time course of the neural response to audiovisual words, measured as the average percent increase in the power of the broadband high-frequency electrical activity (70 to 150 Hz) relative to pre-stimulus baseline, with all responses time-locked to the onset of auditory speech. A superior fit of the additive model would indicate little or no interaction between modalities. A superior fit of the interaction model would indicate subadditivity (if the interaction was negative) or superadditivity (if the interaction was positive).

The additive model decomposed the measured response into two separate time courses, an auditory response and a visual response, under the null hypothesis of no interaction between the modalities. As shown for a single electrode in Fig. 2B, the prediction of the additive model was a poor fit to the measured response (Fig. 2B) with a predicted peak response much greater than the actual peak response (1239% *vs*. 1114% increase from baseline).

The interaction model decomposed the measured response into three separate time courses, an auditory response, a visual response, and their interaction. The interaction model was a better fit to the measured response (Fig. 2C) with similar predicted *vs.* actual peak values (1031% *vs.* 1114%).

To compare the goodness of fit of the two models, we measured the total error over the entire response to auditory speech (−100 to 1000 ms after auditory speech onset). For this electrode, the total error for the additive model was more than twice the error of the interaction model (Fig. 2D; 145% *vs.* 43%).

Next, we fit the additive and interaction deconvolution models to all 33 pSTG electrodes. Averaged across electrodes, the additive model was a poor fit to the measured response with the predicted peak response greater that the actual peak (Fig. 2E; 344% *vs.* 274%). In contrast, the interaction model produced a better fit with similar predicted and actual peak responses (Fig. 2F; 282% *vs.* 274%). To show the results for all individual electrodes, we plotted the total error of both models against each other (Fig. 2G). All of the electrodes were above the line of identity, indicating more error for the additive model. To quantify the model difference across electrodes, a linear mixed-effects model with fixed factor of model type (additive *vs.* interaction, with additive used as the baseline) and random factors of participant and electrode nested within participant was used to compare the models. There was a significant effect of model type, with a parameter estimate of −22±4, *t*(32) = −5.97, p = 10^−6^, demonstrating significantly better fit for the interaction model.

### Nature of the auditory, visual and interaction responses

Examining the modeled auditory, visual and interaction terms allowed us to better understand the multisensory computations in pSTG (Fig. 3A). Averaged across electrodes, the estimated auditory BHA showed a rapid rise from auditory onset (0 ms) to a peak value of 299% at 120 ms; the mean value of the auditory response in the first half-second of the response was 116%. The interaction time series also showed a transient response, but of opposite polarity to the auditory response, with a peak deflection of −79% at 130 ms and an average amplitude of −26% in the first half-second. The visual response showed a different profile, characterized by a more sustained time course without clear peaks. The mean value was 55% in the first half-second of the visual response.

### Heterogeneity across electrodes

There was substantial heterogeneity in the responses of different electrodes. While all electrodes showed a positive response to visual speech, the amplitude of the visual response (averaged of first 500ms) varied between electrodes, from 3% to 280%. The amplitude of the interaction response also varied greatly between electrodes, from −152% to 25%; for 25 out of 33 electrodes, the interaction value was negative, indicating a subadditive interaction (mean value −26, *V* = 79, *p* = 10^−4^, from a non-parametric Wilcoxon test similar to a paired *t*-test). To determine if the heterogeneity in visual and interaction amplitudes were related, we correlated the two values (all reported correlations are Spearman rank correlations to minimize the influence of extreme values). Electrodes with positive visual responses showed negative interaction terms, producing a strong negative correlation between the two quantities, *r(31)* = −0.86, *p* = 10^−6^ (Fig. 3B).

This high correlation could be the result of an uninteresting effect: more responsive electrodes might simply show larger auditory, visual and interaction effects. To determine if this was the case, we performed a partial correlation analysis that predicted the interaction response from the auditory and visual response, electrode by electrode. Taking changes in the auditory response magnitude into consideration, the visual response remained predictive of the interaction, *r(31)* = −0.81, *p* = 10^−7^. In contrast, with visual responses taken into consideration, the size of the auditory response did not predict the size of the interaction, *r(31)* = 0.04, *p* = 0.84. The results of the partial correlation analysis demonstrate that the amplitude of the visual response, but not the auditory response, predicted the size of the interaction.

### Anatomical differences in the response

More posterior electrodes showed a greater visual response, (Fig. 3C; spatial correlation between anterior-to-posterior electrode location and visual response, |*r(29)*| = 0.49, *p* = 0.006) and more negative interaction values (Fig. 3D; |*r(29)|* = 0.55, *p* = 0.002). Linear mixed-effects models were constructed to estimate whether these spatial patterns were consistent across participants. The models found a significant relationship between electrode location and visual response [*X^2^*(1) = 7.2, *p* = 0.007] and interaction size [*X^2^*(1) = 6.3, *p* = 0.01].

As would be expected given that posterior electrodes had stronger interactions, posterior electrodes were poorly fit by the additive model (correlation between A-P location and additive model error minus interaction model error, |*r(29)|* = 0.39, *p* = 0.03).

Across electrodes, the median time of peak interaction was 160 ms. To determine if this value changed along the pSTG, we estimated the time of peak interaction for each individual electrode. More posterior electrodes showed a shorter time-to-peak of the interaction, |*r(29)|* = 0.41, *p* = 0.02).

### Comparison of deconvolved responses with direct measurements

Next, we sought to check the reasonableness of the estimated responses. Qualitatively, the deconvolved responses to auditory and visual speech measured in the present study appeared similar to the direct measurements of responses to auditory-only and visual-only speech in an earlier study (Karas et al., 2019). To quantify the similarity, we took advantage of the fact that one subject with five speech-responsive electrodes located over pSTG participated in both studies. We correlated the magnitude of the BHA power measured with deconvolution (data from the present study, stimuli consisting of audiovisual words with jitter) and BHA power measured directly (data from the earlier study, stimuli consisting of the auditory-only and visual-only words “drive” and “last”, not used in the current study). There was high overall correlation between the measured responses in the previous study and the deconvolution estimates produced here (r^2^ = 0.86 for AV, r^2^ = 0.90 for A, r^2^ = 0.98 for V).

### Exemplar Differences

The present study presented four different audiovisual words at three different asynchronies. To quantify differences between stimulus conditions directly (without deconvolution), a single value for the response to each condition was calculated (mean BHA for first 500ms after auditory onset). As shown in Figure 3-2, there were consistent differences across stimulus exemplars, with the largest response for auditory-advanced words, a smaller response to unedited words, and the smallest response to visual-advanced words. To quantify this observation, the response amplitudes entered into a linear mixed-effects model with a fixed factor of asynchrony (auditory- advanced, unedited, visual advanced) and random factors of stimulus exemplar, participant and electrode nested within participant. The random effect of stimulus exemplar was modeled separately for each level of asynchrony, generating a random intercept and random slope for each exemplar across asynchrony. With this model, there was a significant effect of asynchrony, *X^2^*(2) = 17, *p* = 10^−4^, demonstrating that the effect of asynchrony was not driven by a particular stimulus exemplar.

**Figure 3-2.**
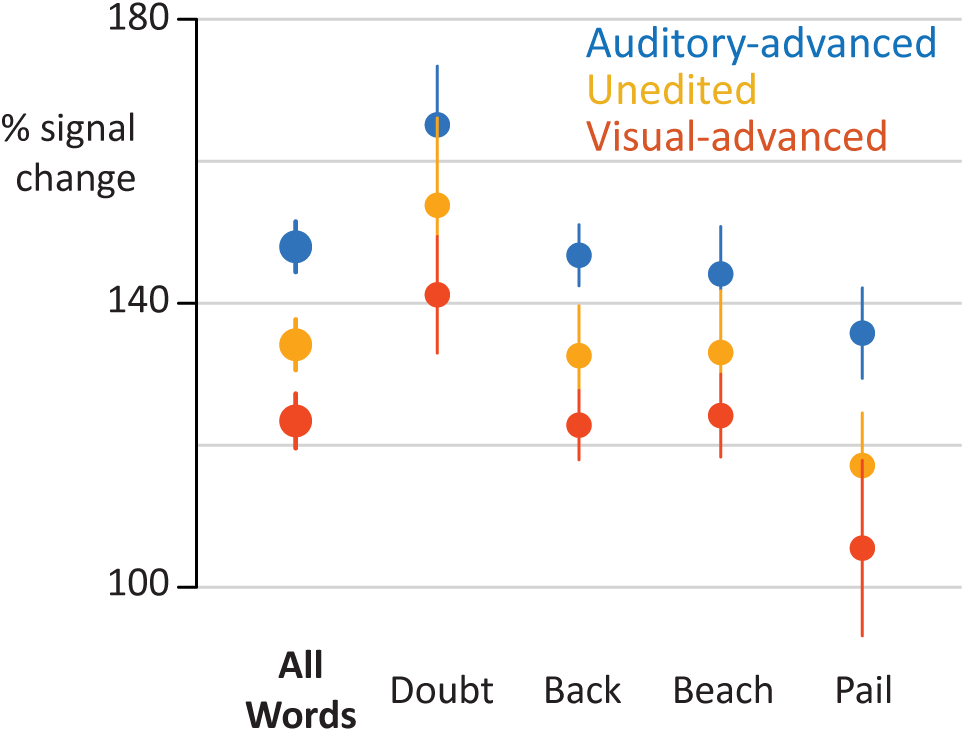
Responses to individual stimulus exemplars (y-axis is percent increase in BHA relative to baseline). Points are colored by asynchrony; error bars are within-subject standard error of the mean.

### Perceptual Reports

After each trial, participants reported whether the auditory and visual speech were perceived as synchronous or asynchronous. Within each asynchrony condition, participants were strongly biased towards one percept or the other. Unedited words were perceived as synchronous on 96.1% of trials; visual-advanced words were rated as synchronous on 81.3% of the trials; auditory-advanced words were rated as synchronous on only 18.7% of the trials (complete behavioral data in Table 1). Within conditions, stimuli were physically identical, but no single condition contained similar numbers of physically-identical trials rated as perceptually different, rendering the data poorly suited for finding perceptually driven responses. Nevertheless, we compared the amplitude of the neural response within each condition to physically identical stimuli that received different perceptual ratings. There were no significant differences in pSTG response as a function of perceptual response for auditory-advanced words (141% for synchronous rating *vs.* 149% for asynchronous rating, two-sample *t*-test *t*(52) = 0.24, *p* = 0.81) or for visual-advanced words (122% for synchronous *vs.* 97% for asynchronous, two-sample t-test *t*(52) = 0.22, *p* = 0.83). There were too few unedited trials rated as asynchronous to perform the comparison.

**Table 1.** Perceptual synchronous/asynchronous task performance. Number of trials for each asynchrony condition for which participants reported a perception of synchronous (”Synch”), asynchronous (”Asynch”) or that were not analyzed (”Excluded”) because there was no response or the neural signal was noisy (greater than ten standard deviations from the mean).

| Subject | Chan<br>count | Unedited |  |  | Auditory-advanced |  |  | Visual-advanced |  |  |
| --- | --- | --- | --- | --- | --- | --- | --- | --- | --- | --- |
|  |  | Synch | Asynch | Excluded | Synch | Asynch | Excluded | Synch | Asynch | Excluded |
| YAI | 6 | 63 | 0 | 1 | 2 | 60 | 2 | 25 | 37 | 2 |
| YAK | 2 | 47 | 0 | 1 | 37 | 7 | 4 | 41 | 3 | 4 |
| YAO | 6 | 59 | 0 | 1 | 0 | 59 | 1 | 57 | 1 | 2 |
| YAQ | 3 | 58 | 0 | 2 | 5 | 52 | 3 | 48 | 10 | 2 |
| YAR | 6 | 56 | 0 | 4 | 0 | 57 | 3 | 58 | 0 | 2 |
| YAS | 6 | 57 | 0 | 3 | 24 | 32 | 4 | 58 | 0 | 2 |
| YCK | 4 | 56 | 1 | 3 | 9 | 49 | 2 | 48 | 11 | 1 |
| Total |  | 396 | 1 | 15 | 77 | 316 | 19 | 335 | 62 | 15 |
| Mean (%) |  | 96.1% | 0.2% | 3.6% | 18.7% | 76.7% | 4.6% | 81.3% | 15.0% | 3.6% |

## Discussion

Using iEEG, we recorded neural responses to audiovisual speech in the human pSTG. While individual stimuli always contained both auditory and visual speech, jittering the onset asynchrony between auditory and visual speech allowed for the time course of the unisensory responses to be estimated using deconvolution. The response to visual speech began at the onset of the talker’s face and showed a flat, sustained response profile. Auditory responses in pSTG were several-fold larger than the visual responses and showed a different response profile, with a transient rise and fall peaking at 130 ms after auditory speech onset. The interaction between auditory and visual speech was subadditive, with a weaker response to audiovisual speech than auditory speech alone, and the degree of subadditivity was correlated with the amplitude of the visual response. Multisensory interactions were strongest in more posterior sections of the pSTG.

### Utility of iEEG deconvolution

Understanding responses to different sensory modalities in isolation is important for understanding the neural computations underlying multisensory integration. However, visual-only speech is an unnatural stimulus that is confusing or even aversive. While we can easily understand auditory-only speech (such as during a phone conversation), visual-only speech is not intelligible. This is a serious confound, as the lack of semantic information might dampen activity in areas of the language network important for semantic processing, or up-regulate areas important for arousal and attention. Deconvolution allowed for the estimation of responses to visual-only speech while avoiding the confounds inherent in presenting it.

It will be important for future studies to investigate how audiovisual interactions differ between the single words used in the present study and the more ethologically relevant stimulus of continuous speech. While auditory-only and audiovisual continuous speech are engaging and easy to understand, participants often report giving up entirely when presented with continuous visual-only speech. By varying the asynchrony, iEEG deconvolution should allow visual and auditory speech components of continuous speech to be separated without this confound. For continuous speech, top-down factors such as expectation and context are also likely to play an important role in modulating pSTG responses (Heald and Nusbaum, 2014; Hickok and Poeppel, 2015; Tuennerhoff and Noppeney, 2016).

The ability of iEEG deconvolution to separate neural responses to auditory and visual speech with only 600 ms of jitter between the two components is due to the high temporal resolution of iEEG. While our study focused on the high-frequency component of the iEEG signal, other studies have measured neural responses to audiovisual speech using ERPs derived from iEEG data (Besle et al., 2008), using MEG (Sohoglu and Davis, 2016), or EEG (Shahin et al., 2012). Deconvolution should be equally applicable to these other techniques since they share the temporal resolution necessary to record millisecond-by-millisecond neuronal responses. Two previous studies applied deconvolution to estimate responses to eye movements in iEEG (Golan et al., 2016) and EEG (Dandekar et al., 2012) datasets.

iEEG deconvolution should also be useful for measuring distinct neural processes underlying a variety of cognitive and perceptual tasks that consist of multiple components that are difficult to separate with subtraction logic, such as mental imagery (Sack et al., 2008) or the encoding, maintenance and retrieval phases of working memory (Baddeley, 2012). Critically, deconvolution allows separation of neural responses to cognitive components that cannot be separated with the traditional Donders subtraction paradigm (Friston et al., 1996).

In fMRI, the BOLD response to even a brief sensory stimulus lasts ~15 seconds. To increase experimental efficiency, in BOLD fMRI rapid event-related designs are used that present stimuli with varying asynchrony on the time scale of seconds (Burock et al., 1998). Both the neural responses and the resulting hemodynamic signals are assumed to sum linearly (Glover, 1999; Henson, 2004). At the faster timescales of iEEG deconvolution, with stimuli jittered by hundreds of milliseconds rather than seconds, we found nonlinear interactions between auditory and visual speech responses: the response in most electrodes was better fit by a model that included an interaction term.

### Classification of Multisensory Computations

A popular schema classifies multisensory neural responses as superadditive, additive, or subadditive by comparing the magnitudes of the responses to unisensory and multisensory stimuli (Stein and Meredith, 1993). To distinguish between additive and non-additive multisensory responses, we fit an additive model and a model that included an interaction term, allowing for super- or sub-additivity. For 25 of 33 electrodes, the interaction term was negative, indicating subadditivity: the response to audiovisual speech was less than the response to auditory-only speech.

These results are consistent with studies recording brain responses from the scalp surface using EEG. In an important early study, Besle and colleagues studied responses to A, V and AV syllables and found evidence for suppressive audiovisual integration (Besle et al., 2004). With similar stimuli, van Wassenhove and colleagues (van Wassenhove et al., 2005) found that the N100/P200 event related potential (ERP) was weaker for audiovisual than auditory-only speech, consistent with the present results. Subadditivity for audiovisual speech has also been demonstrated for the N200 and N400 components of the speech ERP (Besle et al., 2004; Pilling, 2009; Ganesh et al., 2014; Paris et al., 2017). Applying the additivity classification scheme to BOLD fMRI data poses methodological difficulties (Beauchamp, 2005; Laurienti et al., 2005) and may explain early reports of superadditivity in pSTG (Calvert et al., 1999; Calvert et al., 2000). Later BOLD fMRI studies typically report subadditivity (Wright et al., 2003; Nath and Beauchamp, 2011; Okada et al., 2013) although see (Werner and Noppeney, 2010).

Recording with intracranial EEG in auditory cortex, Besle and colleagues found evidence for visually-driven reductions in responses to auditory speech (Besle et al., 2008), although it is important to note that the event-related potentials (ERPs) used as a measure of neural activity by Besle and colleagues differs from the broadband high-frequency power (BHA) measured in the present study. In general, BHA amplitude correlates with the rate of action potential firing by nearby neurons (Ray et al., 2008; Ray and Maunsell, 2011) although it may also include contributions from dendritic processing (Leszczynski et al., 2019) while ERPs are thought to reflect summed synaptic potentials (Buzsaki et al., 2012).

Besle and colleagues (Besle et al., 2008) suggested that “preprocessing of the visual syllable would result in engaging less auditory resources from the auditory cortex”, resulting in the observed reduction in auditory responses compared with visual responses. Building on this suggestion, in a recent paper we suggested that this visual preprocessing could selectively inhibit populations of neurons response to auditory phonemes incompatible with the observed visual mouth shape (Karas et al., 2019). The model incorporates evidence that the pSTG contains neural populations that represent specific phonemes (Formisano et al., 2008; Mesgarani et al., 2014; Hamilton et al., 2018) and that visual information influences processing in auditory cortex (Calvert et al., 1997; Pekkola et al., 2005; Besle et al., 2008; Kayser et al., 2008; Zion Golumbic et al., 2013a; Zion Golumbic et al., 2013b; Rhone et al., 2016; Megevand et al., 2018; Ferraro et al., 2019). Reduced responses in pSTG to audiovisual speech may reflect more efficient processing, with less neural resources required to decode the speech. In turn, increased processing efficiency in pSTG could explain behavioral observations that audiovisual speech perception is faster, more accurate, and less effortful than auditory-only speech perception (Moradi et al., 2013). Support for a selective inhibition model is also provided by an elegant EEG experiment (Shahin et al., 2018). Using incongruent audiovisual speech pairings, visual speech modulated both the perception of an incongruent auditory syllable and rendered the evoked N100 component of the ERP similar to the congruent auditory condition. Suppressive interactions in higher-order cortex may be a general property of complex computations, as they have also been observed in other systems, including visuomotor integration in the frontal eye fields (Mirpour et al., 2018).

Although pSTG was classified by Brodmann as a single anatomical compartment (area 22) there is increasing evidence that anterior and posterior pSTG are functionally distinct (Ozker et al., 2017; Hamilton et al., 2018; Ozker et al., 2018b). Even within parts of posterior pSTG that are visually responsive, there is evidence that the cortex is divided into subregions that respond to the visual mouth movements that make up visual speech and subregions that respond to viewed eye movements (Zhu and Beauchamp, 2017; Rennig and Beauchamp, 2018). The present study lends credence to the idea that there is substantial heterogeneity in this region, with multisensory interactions concentrated in the posterior portions of pSTG.

## Acknowledgments

This research was supported by NIH R01NS065395 and R25NS070694.

